# Large-scale characterisation of the pregnancy vaginal microbiome and sialidase activity in a low-risk Chinese population

**DOI:** 10.1101/2021.06.02.446856

**Authors:** Sherrianne Ng, Muxuan Chen, Samit Kundu, Xuefei Wang, Zuyi Zhou, Zhongdaixi Zheng, Wei Qing, Huafang Sheng, Yan Wang, Yan He, Phillip R. Bennett, David A. MacIntyre, Hongwei Zhou

**Author notes:** Corresponding authors: D.A.M.,; H.Z. These authors contributed equally to this study (Sherrianne Ng, Muxuan Chen).

## Abstract

Vaginal microbiota-host interactions are linked to preterm birth (PTB), which continues to be the primary cause of global childhood mortality. Despite the majority of PTB occuring in Asia, studies of the pregnancy vaginal microbiota are largely limited to Northern American and European populations. Here, we characterised the vaginal microbiome of 2689 pregnant Chinese women using metataxonomics and in a subset (n=823), the relationship between vaginal microbiota composition, sialidase activity and leukocyte presence and pregnancy outcomes. Vaginal microbiota were most frequently dominated by *Lactobacillus crispatus* or *L. iners*, with the latter associated with vaginal leukocyte presence. Women with high sialidase activity were enriched for bacterial vaginosis-associated genera including *Gardnerella, Atopobium* and *Prevotella*. Vaginal microbiota composition, high sialidase activity and/or leukocyte presence was not associated with PTB risk suggesting underlying differences in the vaginal microbiota and/or host immune responses of Chinese women, possibly accounting for low PTB rates in this population.

**Importance:** Specific vaginal microorganisms or ‘vaginal microbiota’, are associated with preterm birth, which is the primary cause of death in children under 5yrs of age worldwide. Despite most preterm births occuring in Asia, almost all studies of the pregnancy vaginal microbiota have been limited to Northern American and European women. Here, we studied the vaginal microbiota in a large cohort of 2689 pregnant Chinese women and showed that it was most frequently dominated by *Lactobacillus crispatus* or *L. iners*. The latter was associated with leukocyte infiltration of vaginal secretions. Women with high activity of the enzyme sialidase, were frequently colonised by species associated with the common condition bacterial vaginosis, including *Gardnerella, Atopobium* and *Prevotella* species. Vaginal microbiota, high sialidase activity and/or leukocyte presence was not associated with preterm birth risk indicating differences in the microbe-host immune responses of Chinese women, possibly explaining why preterm birth rates in this population are low.

## Introduction

There is substantial evidence implicating the pregnancy vaginal microbiota in shaping maternal and neonatal health outcomes (1–3). Dominance of the vaginal niche by commensal *Lactobacillus* species is often considered “optimal” due to their ability to prevent pathogen colonisation through competitive exclusion, in part achieved through the production of antimicrobial compounds and production of lactic acid (4, 5). Recent studies have highlighted *L. crispatus* dominance as being protective against preterm birth (PTB)(2,6–9) and neonatal sepsis following preterm prelabour rupture of membranes (1). By contrast, colonisation by *L.iners* (7,10–12) or *Lactobacillus* species depleted, high diversity compositions are associated with an increased risk of PTB (2,7,9,10,13–15). Despite the majority of PTBs (60%) occurring in Asia (16), molecular based characterisation of the vaginal microbiota in pregnancy and its relationship with PTB has largely been restricted to Northern American and European populations. Moreover, ethnicity is now recognised as a potential confounder of the relationship between the vaginal microbiome and PTB, particularly between Caucasian and women of African-descent in North American or European populations (2,9,10,17).

The Amsel criteria (18) and Nugent scoring system (19) are commonly used to diagnose Bacterial Vaginosis (BV), a common condition characterised by a loss of vaginal lactobacilli and overgrowth of anaerobes, which is associated with a two-fold increased risk of PTB (20–23). However, both techniques require laboratory access and microscopy, and are inherently subjective. Enzymatic based assays for rapid BV diagnosis may offer an objective, point-of-care alternative to BV diagnosis in clinical settings, including during pregnancy (24, 25). These assays often work by measuring microbial sialidase (neuraminidase), produced by BV-associated bacteria such as *Gardnerella vaginalis*, which removes sialic acid from sialoglycoconjugates including those on the surface of vaginal epithelial cells, providing a nutrient source and exposing glycan-binding sites for bacterial adhesion. Additionally, sialidase is thought to mediate biofilm formation and the establishment of sub-optimal vaginal microbiota compositions (26–28). High vaginal sialidase levels have previously been associated with an increased risk of PTB (29) and with failure of cervical cerclage (30), a procedure used to reinforce the cervical opening in women at risk of preterm delivery due to cervical shortening. Sialidase-producing taxa associated with BV have also been implicated in chorioamnionitis, a risk factor for PTB that is characterised by inflammation of the fetal membranes (31). Although BV is not classified as an inflammatory syndrome, the disease has been associated with the presence of vaginal leukocytes, which have been purported to offer predictive value in identifying upper reproductive tract infections (32–34). Quantification of leukocyte counts using vaginal wet mount microscopy could therefore represent an easily accessible and cost-effective method to determine cervicovaginal inflammation (35).

In this study, we characterised the bacterial component of the vaginal microbiome in 2689 Chinese women sampled at mid-pregnancy and in a subset (n=823), explored the relationship between vaginal microbiota composition, sialidase activity and leukocyte presence with risk of PTB.

## Results

### Study population

A total of 2796 women, with a median maternal age of 29 years (18 – 47 years), met the study inclusion criteria and were recruited between November 2015 to December 2018. The mean time of sampling was 16^+4^ weeks^+days^ gestation (range 11 – 40^+5^). Of these, 1405 delivered at the same hospital and maternal and neonatal outcomes data were obtained. The median gestation at delivery was 39^+3^ (20^+3^ – 41^+5^) weeks^+days^. The PTB (<37 weeks) rate in the cohort was 5.4% (76/1405)(Data set S1). The remaining 1391 women delivered elsewhere and due to data protection, pregnancy outcome data was not available. There was no significant difference in gestation of sampling between women with and without available outcome data (Mann-Whitney Test, p>0.05; Fig. S1).

### The pregnancy vaginal microbiota of Chinese women

The 2796 sequenced vaginal samples generated a total of 58,582,840 reads with a mean read count of 20,952 per sample. Of these, 2689 passed library size and microbiome classification criteria (Fig. S1). After removal of kit and reagent contaminants, a total of 82 taxa were detected and included in subsequent analyses. Vaginal microbiota profiles were classified into 19 groups based upon the dominant (>30% relative abundance) taxa observed within each sample (2). At the genus level, the majority of samples were dominated by *Lactobacillus* (2311/2689, 85.94%), which were predominately *L. crispatus* (1077/2689, 40.05%) or *L. iners* (966/2689, 35.92%) dominated (Fig. 1a). The median relative abundance of *L. crispatus* and *L. iners* in these samples was 96.32% and 96.21% respectively and the alpha diversity as measured by the Shannon Index was similar between the two groups (median, 0.22 and 0.24) (Table S1). Conversely, microbiota dominated by BV-associated species including *Gardnerella* spp. (225/2689, 8.37%), *Prevotella* spp. (25/2689, 0.93%) and *Atopobium* spp. (26/2689, 0.97%) had lower relative abundances (<70%) and higher alpha diversity (Shannon Index >0.85) compared to samples classified as *L. crispatus* or *L. iners* dominated (Shannon Index <0.25, Kruskall-Wallis Test, p<0.001) (Fig. 1a and Table S1). When these dominant taxa profiles were grouped into Vaginal Microbiome Groups (VMG) consistent with previously reported “community state types” (36) or “vagitypes” (2), the most prevalent was VMG I (1077/2689, 40.05%), followed by VMG III (966/2689, 35.92%), VMG IV-A (287/2689, 10.67%), VMG II (138/2689, 5.13%), VMG V (97/2689, 3.61%) and IV-B (31/2689, 1.15%). Groups dominated by other *Lactobacillus* species or *Bifidobacterium* species were classified into VMG VI (33/2689, 1.23%) and VII (60/2689, 2.23%)(Fig. 1b and Table S2), respectively. The proportions of VMGs in women with and without available pregnancy outcome data (Fig. S1) were similar (Fisher’s Exact Test, p=0.22).

**Figure 1.**
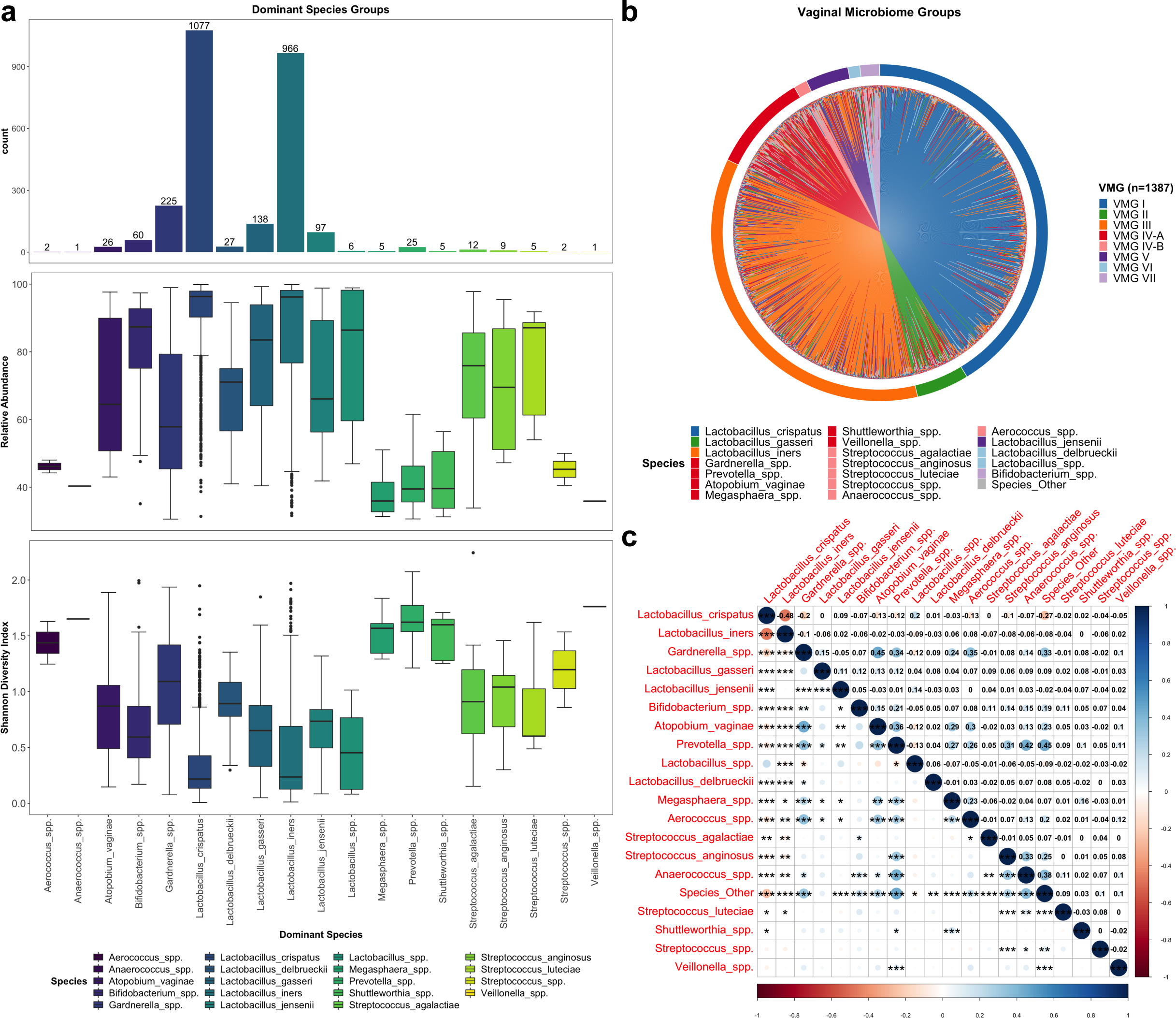
Vaginal microbiome composition and structure in low-risk Chinese pregnancy. (a) Bar charts display number of samples within each dominant species group with box plots presenting median and interquartile range for relative abundance of species and Shannon diversity index of each dominant species group. (b) Microbiota profiles of women with outcome data classified by vaginal microbiome groups (VMG). Colour of species denotes the VMG group classification with VMG I (*L. crispatus* dominated, blue), II (*L. gasseri* dominated, green), III (*L. iners* dominated, orange), IV-A (bacterial vaginosis associated, red), IV-B (pathobionts, pink), V (*L. jensenii* dominated, purple), VI (Other *Lactobacillus*, light blue), and VII (*Bifidobacterium* dominated, light purple). (c) Correlation plot for dominant species showing correlation strength (circles), relationship (positive, blue; negative, red) and significance (***, p<0.001; **, p<0.01; *, p<0.05).

Relative abundance of *L. crispatus* was found to be negatively correlated to the majority of vaginal taxa including *L. iners* (r=-0.48, p<0.001). *L. iners* was also negatively correlated to the majority of other vaginal taxa except *Megasphaera* spp. (r=0.06, p<0.05) and *Aerococcus* spp. (r=0.08, p<0.01), where weak positive correlations were observed. Several BV-associated pathogens were positively correlated with each other including *Gardnerella* spp., with *Atopobium vaginae* (r=0.45, p<0.001), *Prevotella* spp. (r=0.34, p<0.001), *Aerococcus* spp. (r=0.35, p<0.001) and Other Species (r=0.33, p<0.001); *Atopobium vaginae* with *Prevotella* spp. (r=0.36, p<0.001), *Megasphaera* spp. (r=0.29, p<0.01), and *Aerococcus* spp (r=0.30, p<0.001); and *Prevotella* spp. with the pathobionts *Streptococcus anginosus* (r=0.31, p<0.001) and *Anaerococcus* spp. (r=0.42, p<0.001) as well as Other Species (r=0.45, p<0.001)(Fig. 1c).

### Sialidase activity and vaginal microbiota structure in pregnancy

High vaginal sialidase activity was strongly associated with increased bacterial alpha diversity (n=36; median Shannon index 0.93 (0.09 – 1.79)) compared to women with low vaginal sialidase activity (n=787; median Shannon index 0.32 (0.01 – 2.24), Mann-Whitney Test, p<0.001)(Fig. 2a). Women with high sialidase activity had significantly higher prevalence of *Lactobacillus* species depleted microbiota (30/36, 83.33% versus 183/787, 23.25%; Fisher’s Exact Test, p= 2.089e-13) and VMG IV-A (22/36, 61.11% versus 58/787, 7.37%; Fisher’s Exact Test, p=0.0005)(Fig. 2b and Fig. S3). VMG I (2/36, 5.56% versus 323/787, 41.04%) and II (2/36, 5.56% versus 48/787, 6.10%) were rarely observed in women with high sialidase activity whereas prevalence of VMG III was similar in women with high and low sialidase activity (10/36, 27.78% versus 288/787, 36.59%). Samples classified as VMG IV-B (11/787, 1.40%), V (35/787, 4.45%), VI (9/787, 1.14%) and VII (15/787, 1.91%) were only associated with low sialidase activity. Differential abundance analysis using LDA and LefSe of samples with high and low sialidase activity confirmed *L. crispatus* and *L. jensenii* as being enriched in low sialidase activity samples whereas high activity was characterised by enrichment with BV-associated genera including *Gardnerella*, *Atopobium*, *Prevotella* and *Megasphaera* (Fig. 2c). In these women, higher total relative abundance of BV-type pathogens (*Gardnerella spp.*, *Atopobium vaginae*, *Prevotella spp.* and *Megasphaera spp.*) was observed compared to women with low sialidase activity (median, minimum – maximum; 18.83 (0.00 – 62.15) v 0.64 (0.00 – 59.55), Mann-Whitney Test, p<0.01)(Fig. 2d and Table S3). Using metataxonomics-defined *Lactobacillus* depleted (positive) and *Lactobacillus* dominated (negative) as the reference test, sialidase activity in this cohort was found to have low sensitivity (14.08%), high specificity (99.02%), moderate positive predictive value (PPV, 71.65%) and high negative predictive value (NPV, 86.72%)(Fig. 2e).

**Figure 2.**
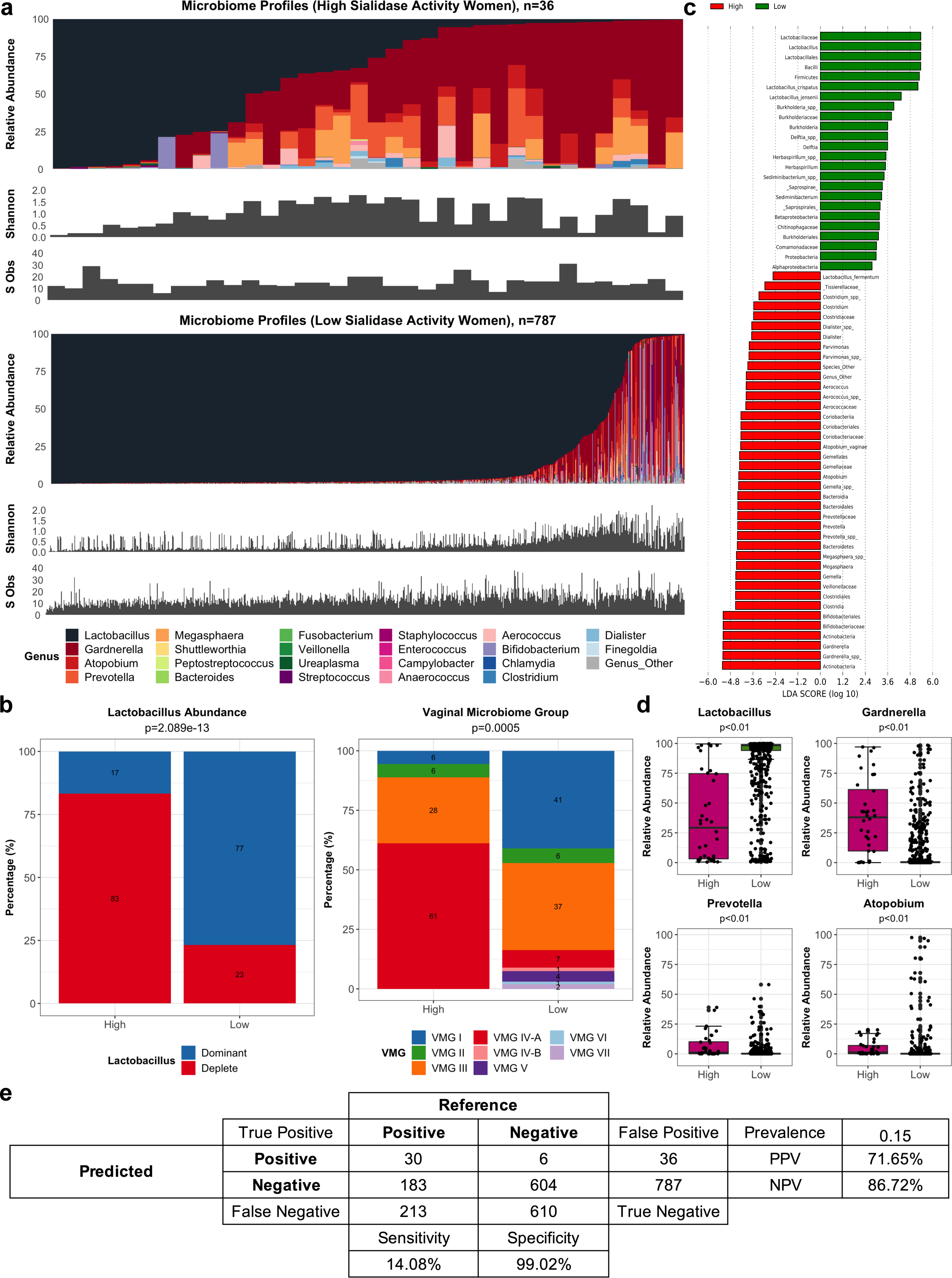
Relationship between sialidase activity and vaginal microbiota composition. (a) Microbiota composition, diversity (Shannon Index) and richness (species observed) for all women with high (n=36) and low (n=787) sialidase activity. (b) Proportion of *Lactobacillus* abundance groups (*Lactobacillus* dominant and *Lactobacillus* deplete) and vaginal microbiome groups (VMG) for women with high or low sialidase activity. (c) LDA showing effect size of differentially abundant taxa associated with microbiota of women with high (red) or low (green) sialidase activity. (d) Comparison of relative abundance between main BV-associated bacteria identified as significantly different (Mann-Whitney Test, p<0.05) between women with high and low sialidase activity (see Supplementary Table 3). (e) Confusion matrix with *Lactobacillus* abundance (dominant as positive/deplete as negative) as the reference test and sialidase result (High/Low) as the predicted test. The sensitivity, specificity, positive predictive value (PPV) and negative predictive value (NPV) were calculated.

### Leukocyte presence and vaginal microbiota composition during pregnancy

Leukocyte +ve vaginal wet mounts were associated with a small but significantly higher proportion of *Lactobacillus* dominated microbiomes (441/574, 76.83%) compared to women with leukocyte -ve vaginal wet mounts (169/249, 67.87%; Fisher’s Exact Test, p=0.01). This relationship was driven by significantly higher VMG III prevalence in these women (Fisher’s Exact Test, p=0.02)(Fig. 3a). This was further confirmed by comparing relative abundance levels of *Lactobacillus* species between Leukocyte +ve and -ve samples (Mann-Whitney Test, p<0.001)(Fig. 3b). Similar results were observed when using LDA and LefSe analysis to identify differential taxa between leukocyte +ve and leukocyte -ve wet mounts with *L. iners* having the strongest effect size of the differentially abundant features identified (Fig. 3c).

**Figure 3.**
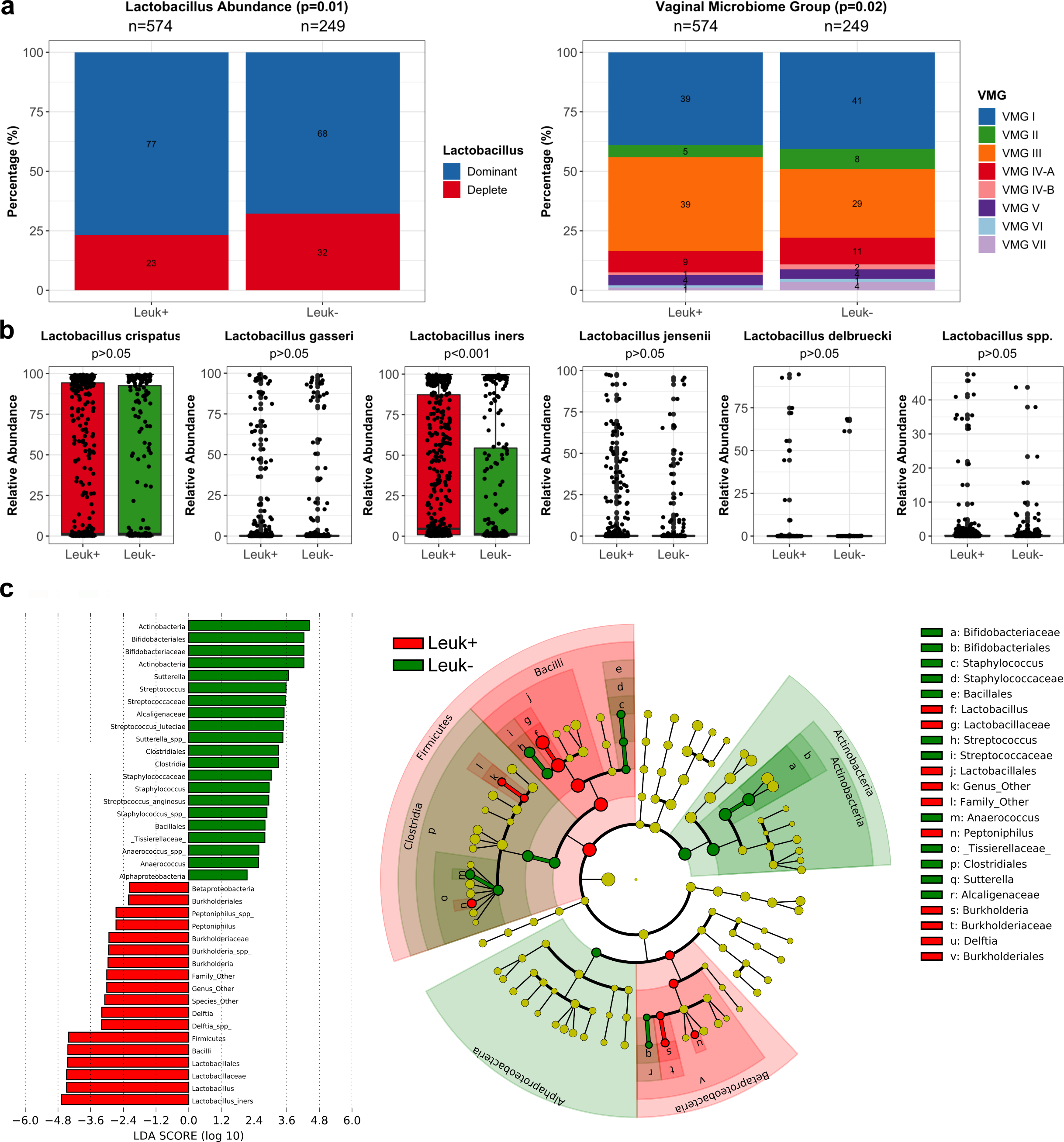
Vaginal microbiome composition and leukocyte presence in vaginal secretions. **(a)** Stacked bar of showing vaginal microbiome groups (VMG) of women with leukocyte presence (Leuk+) or absence (Leuk-). **(b)** Boxplots showing relative abundance of *L. crispatus*, *L. gasseri*, *L. iners*, *L. jenseni*, *L. delbruecki* and *Lactobacillus* spp. in women with leukocyte presence (Leuk+, red) and absence (Leuk-, green). **(c)** LDA of differentially abundant taxa and cladogram displaying bacterial clades and nodes identified as differentialy abundance from LDA analysis in vaginal microbiota of women with leukocyte presence (Leuk+, red) and absence (Leuk-, green). Statistical significance of categorical variables based on Fisher’s Exact Test and continuous variables based on Mann-Whitney Test.

### Vaginal microbiota, sialidase activity and leukocyte presence are not associated with risk of PTB and chorioamnionitis in pregnant Chinese women

In women with available outcome data (n=1387), the proportion of *Lactobacillus* dominated or depleted vaginal microbiota (p=0.57; Fisher’s Exact test) and VMGs (p=0.05; Fisher’s Exact test) were similar between women delivering preterm or term. Likewise, proportions of high or low sialidase activity (p=0.45; Fisher’s Exact test) and leukocyte presence/absence (p=0.74; Fisher’s Exact test) were comparable between women subsequently experiencing preterm or term deliveries (Fig. 4a). Apart from women who had *Lactobacillus* dominated microbiomes, high sialidase activity and leukocyte absence which had insufficient sample size (n=2), no significant difference was observed in birth gestation based on *Lactobacillus* abundance, VMG prevalence, sialidase activity and leukocyte wet mount results (Kruskal-Wallis Test, p>0.05)(Fig. 4b and Table S4). Logistic regression results similarly showed vaginal microbiome *Lactobacillus* composition (p=0.83), sialidase activity (p=0.45) and leukocyte presence or absence (p=0.70) did not significantly contribute to birth outcome. Finally, no relationships were observed between vaginal microbiota composition, sialidase activity and/or leukocyte presence and chorioamnionitis (Fig. S4).

**Figure 4.**
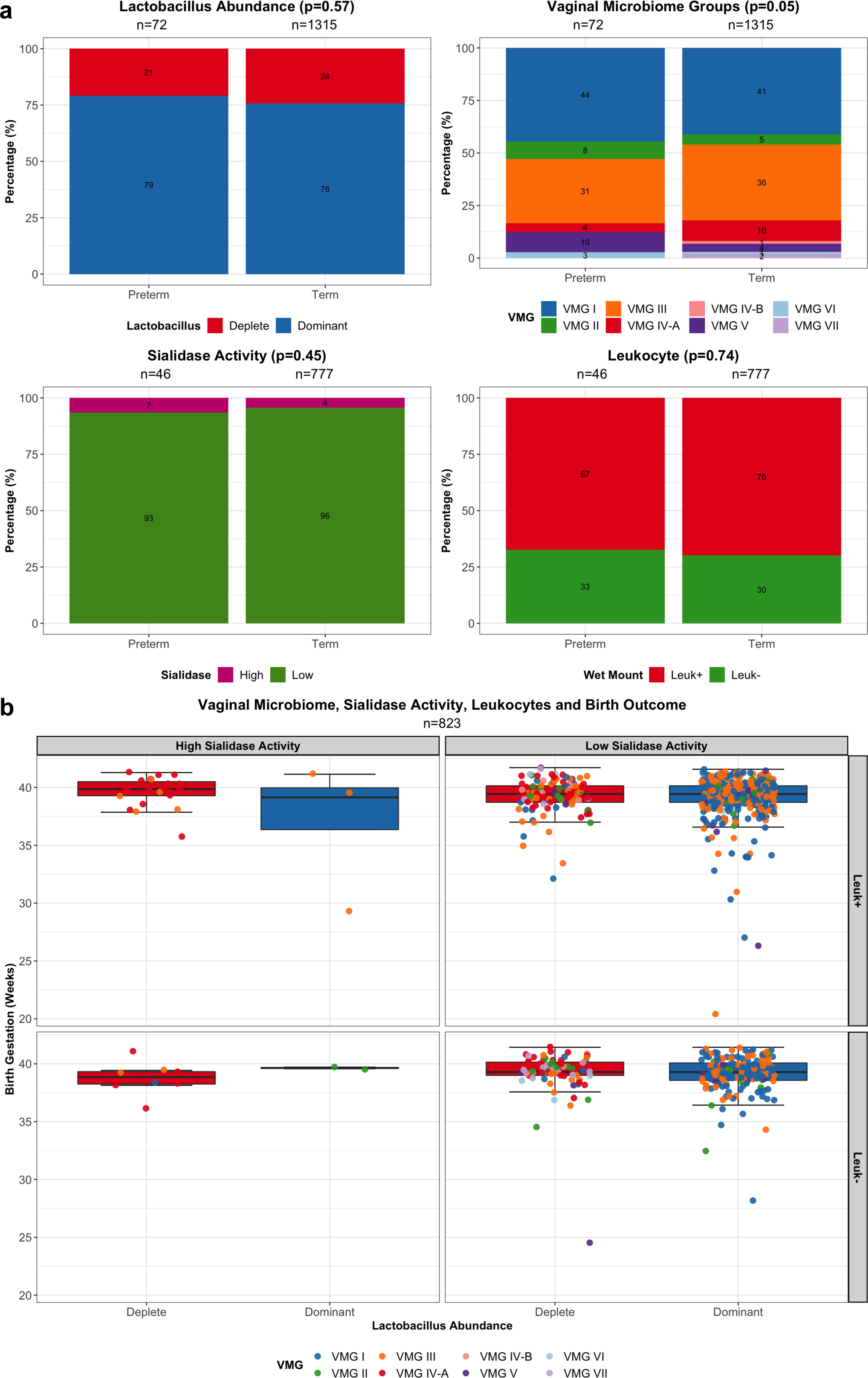
Association between vaginal microbiota composition, sialidase activity and leukocyte presence and birth outcomes. **(a)** Stacked bar plots showing proportion of *Lactobacillus* abundance (deplete, red; or dominant, blue), vaginal microbiome groups (VMG), sialidase activity (high, pink; or low, red) and leukocyte presence (Leuk+, red) or absence (Leuk-, green) in preterm and term groups. Statistical significance based on Fisher’s Exact Test. **(b)** Boxplots showing birth gestation in weeks grouped by *Lactobacillus* abundance, sialidase activity and leukocyte presence or absence with dot points colored by VMG. No significant difference between boxplots with n≥3 based on Kruskal-Wallis Test, p>0.05.

## Discussion

In this large cross-sectional study of 2689 pregnant Chinese women, the vaginal microbiota in mid-pregnancy was characterised by *Lactobacillus* spp. dominance and low diversity. This pattern is consistent with previous descriptions of the healthy mid-pregnancy vaginal microbiome in Caucasian, African and Asian women from European and Northern American populations (6,10,13,17,37). Similar results have also been reported in small cohort studies of women of Chinese and Indian descent outside of western geographical regions (38, 39). One of these studies examined the vaginal microbiota of 113 Chinese women and reported *L. crispatus* dominance in 45.1% and *L. iners* dominance in 31.9% of women sampled at a mean gestation of 17 weeks (38). In our cohort, *L. crispatus* (40%) and *L. iners* (36%) dominated vaginal microbiota were also the most frequently observed profiles. In previous work, we have also observed that the most prevalent vaginal microbiota community compositions in European Caucasian women are those dominated by *L. crispatus* (50%) followed by *L. iners* (25%), whereas the inverse was seen for women self identifying as Asian and Black (*L. iners,* 48% and 50% v *L. crispatu*s, 26% and 20%, respectively)(6). Similarly, higher frequency of *L. iners* dominated vaginal microbiota has been reported in Indian (39), Karen and Burman (40), Hispanic and African American pregnant women (9,10,17). Our results therefore provide further evidence that ethnic and/or geographical differences are a major source of variation in the underlying structure and composition of the vaginal microbiome during pregnancy.

Correlation analyses in our patient cohort showed that *L. crispatus* was negatively correlated with almost all other vaginal taxa, highlighting its well-described exclusionary behaviour in the vaginal niche that appears to be broadly consistent across different ethnic groups (6,13,17,41). However, we also observed a strong negative correlation between *L. iners* and most other vaginal taxa. This was surprising given that *L.iners* has been shown to co-occur with various BV-taxa in other ethnic groups including Caucasian and African American women (9, 42). The highly negative correlation observed between *L. iners* and *L. crispatus* might be attributable to overlapping ecological functions in the vagina (5, 43). Compared to *L*.

*crispatus*, *L. iners* has a substantially smaller genome, thought to be indicative of a symbiotic and/or parasitic role in the vaginal niche (5,43,44). Similar to *G. vaginalis*, *L. iners* can produce pore-forming toxins such as inerolysin and vaginolysin, which can lead to lysis of host cells and release of carbon sources, particularly during times of nutrient scarcity (5,43,44). This behaviour may explain the observed positive relationship between *L.iners* and leukocyte presence in our study cohort. Consistent with these findings, a recent study of 83 healthy Chinese pregnant women reported a positive association between leukocyte esterase concentrations and increased *L. iners* levels determined using quantitative real-time PCR (45). These women were also reported to have increased presence of pus cells and other non-Lactobacillus morphotypes observed under microscopy. *In vitro* experiments using human vaginal epithelial cells have also demonstrated that *L. iners* stimulates increased pro-inflammatory cytokine production compared to *L. crispatus* (46, 47). Despite these findings, we did not observe any relationship between *L. iners* dominance or any vaginal microbiota profile and subsequent risk of PTB in this cohort. This is in contrast to previous studies of predominantly Caucasian women by ourselves and others that have reported a relationship between *L.iners* and increased risk of PTB or cervical shortening, which is clinically used as a marker of PTB risk (6,7,12,48). Our data instead indicates that like Hispanic and African American women, *L. iners* does not appear to be risk factor for PTB in Chinese populations.

In our sub-cohort analysis, high vaginal sialidase activity was strongly associated with *Lactobacillus* depleted microbiota enriched with BV-associated taxa including *Gardnerella*, *Prevotella*, *Atopobium*, and *Megasphaera* species. This finding is consistent with previous studies in both pregnant and non-pregnant women where the production of sialidase by BV-associated pathogens, such as *Gardnerella* spp. and *Prevotella* spp., is thought to be important for pathogenesis (26,49,50). In this context, desialylation of glycolipids and/or glycoproteins (e.g. immunoglobulins, cytokines, cellular receptors, mucins and antimicrobial molecules) can decrease ability of host defence responses to recognise and bind to microbes whilst increasing bacterial adherence, invasion and tissue breakdown (26,29,50,51). Interestingly, high sialidase activity was observed in some *Lactobacillus* dominated vaginal microbiota. These often contained low relative aundance of BV-associated taxa indicating that perhaps even low levels of sialidase-producing bacteria may be sufficient to produce high sialidase activity. It is also possible that sialidase was produced by other microorganisms including viruses and fungi that were not assessed in our study (52, 53). In contrast to these findings, a proportion of samples (23%) harbouring *Lactobacillus* spp. depleted vaginal microbiota enriched with BV-associated taxa did not have high sialidase activity. This was supported by our findings indicating sialidase activity has high specificity (99%) but low sensitivity (14%) in predicting high-diversity, *Lactobacillus* spp. depleted vaginal microbiota. Low sialidase activity in these samples could be due to several factors: 1) the production of sialidase by BV-taxa, including *Gardnerella* spp., is strain-dependent (26,29,30); 2) binding of sialidase to the BV test kit substrate may be influenced by vaginal pH (25, 54); 3) the amount of sialidase produced by BV-taxa maybe insufficient to reach the detection limit of the BV test kit; and/or 4) the bacterial load of BV-taxa, which was not quantified in our study, may be inadequate to produce sufficient sialidase for detection by the BV test kit. Overall, our findings suggest that high sialidase activity is significantly associated with BV-taxa but cannot be used as an accurate proxy for high-diversity, *Lactobacillus* spp. depleted vaginal microbiota in pregnant women.

Elevated sialidase activity has previously been associated with spontaneous PTB and late miscarriage in a North American study of 1806 pregnant women when measured at 12 weeks gestation (29), and with early PTB and low birth weight in a mid-trimester study of 579 Danish women (55). Additionally, in a recent study of 85 Chinese women, sialidase activity was associated with subsequent cervical cerclage failure, a risk factor for PTB (30). However, in our study, no significant association was found between elevated sialidase activity and PTB. Our findings are similar to a previous nested case-control study of 126 pregnant women, where elevated sialidase activity measured at mid-pregnancy was not associated with PTB risk (49). The observed differences between studies could be due to different methods for measuring sialidase activity where a single cut-off was used rather than a quantitative measurement. An association between vaginal leukocytes and preterm labour or risk factors of PTB including reproductive tract infections, histologic chorioamnionitis and inflammatory cytokines have also previously been reported (32,33,35,56–58). Here, we classified leukocyte presence based on vaginal wet mount microscopy, which has been shown to be useful for identification of cervicovaginal inflammation in pregnant women (35). Logistic regression analyses indicated that neither leukocyte presence, sialidase activity nor vaginal microbiota contributed to preterm risk in our study cohort. The rate of PTB in our study population was 5.4% which is slightly lower than the expected background rate in China (7.2%)(59). However, the rate of PTB in China is considerably less than the global average of ∼10% (60). Taken together, our findings suggests that Chinese women may have underlying differences in the vaginal microbiota and/or host immune responses that potentially contribute to lower rates of PTB. A limitation of our study is the lack of longitudinal sampling of the patient cohort, which would have facilitated deeper analysis of vaginal microbiota dynamics across pregnancy and its relationship with sialidase activity, leukocyte presence and birth outcomes.

In conclusion, our study provides new insight into the vaginal microbial composition and structure of Chinese pregnant women. Although sialidase activity was predictive of high diversity vaginal microbiota compositions, neither were associated with increased risk of PTB. *Lactobacillus* spp. depleted vaginal microbiomes with high sialidase activity and leukocyte presence are not associated with higher risk of PTB in Chinese women. Our results provide further evidence that ethnicity is an important determinant of microbiota-host interactions during pregnancy and highlight the need for further investigations into the mechanisms underpinning these relationships.

## Methods

### Study design

Pregnant women at their first prenatal visit were recruited from Nanfang Hospital of Southern Medical University in Guangzhou, China, from January 2015 to December 2018. Written, informed consent was obtained from all participants. Women who received antibiotics, prebiotics or probiotics within 30 days prior to vaginal swab collection and/or who had sexual activity within 48 hours of sample collection were excluded. Metadata collected included maternal age, gestation at delivery (days) and chorioamnionitis. This prospective study was reviewed and approved ([2013] EC (100)) by the Ethical Committee of Nanfang Hospital, Southern Medical University, Guangzhou, China.

### Sample collection and processing

Vaginal samples were collected using a sterile swab that was inserted into the vaginal posterior fornix, before being gently rotated 360° for approximately 20 rotations prior to removal. All swabs were placed immediately on ice and were then stored at -20°C within 1 hour of collection before being transferred to -80°C within 24 h for long-term storage. For DNA extraction, swabs were immersed and vigorously mixed in 500 μL of sterile water before the solution was transferred to a clean 2 mL centrifuge tube. The sample was then vortexed for 5 sec before being centrifuged at 13,800 *g* for 10 min. The supernatant was removed, and the pellet retained for DNA extraction. For PCR negative controls (n=82), 2 μL of diethylpyrocarbonate (DEPC) water was used as non-template control. For DNA extraction kit negative controls (n=21), 500 μL of sterile water was used instead of test samples for the DNA extraction process.

### DNA extraction and 16S rRNA (V4) amplicon sequencing

DNA was extracted from vaginal swabs using either BioTeke bacterial DNA extraction kit (BioTeke Corporation, Cat #DP7001) per manufacturer’s instructions (n=1176) or Bioeasy bacterial DNA extraction kit (Shenzhen Bioeasy Biotechnology Co. Ltd., Cat #YRMBN7001) per described for Thermo Scientific^TM^ KingFisher^TM^ Flex Purification System (n=1620). The V4 region of bacterial 16S rRNA was amplified using barcoded V4F 5’ GTGCCAGCMGCCGCGGTAA 3’ forward and V4R 5’ CTACCNGGGTATCTAAT 3’ reverse primers. The PCR condition included an initial denaturation step at 94°C for 5 min, 30 cycles of 94°C for 30 sec, 52°C for 30 sec, 72°C for 45 sec and a final extension step at 72°C for 5 min. Each 20 μl reaction volume consisted of 10 μl of AceQqPCR SYBR Green Master Mix (Vazyme, Nanjing, China), 0.4 μL prime V4F, 0.4 μL primer V4R, 0.4 μL ROX Reference Dye 2, 2 μL template DNA, 8.8 μL sterilized distilled water. Equimolar amplicon suspensions were combined and subjected to paired-end 101 bp sequencing on an Illumina HiSeq sequencer at the Beijing Genomics Institute (BGI; Beijing, China).

### Sialidase enzyme activity detection

A total of 823 women had an additional vaginal swab collected for detection of sialidase enzyme activity. Sialidase enzyme activity was measured as per manufacturer’s instructions using a single-enzyme BV kit based on the colorimetric method (Zhuhai DL Biotech. Co. Ltd). Briefly, each swab sample was immersed into a BV test bottle solution containing 5- bromo-4-chloro-3-indolyl-α-D-N-acetylneuraminic acid (BCIN) as a substrate (61). The BCIN substrate hydrolyses and reacts with the added 1 – 2 drops of BV chromogenic solution containing hydroxide and potassium acetate, resulting in a colour reaction that was measured using the BV-10 analyser (Zhuhai DL Biotech. Co. Ltd). The resulting sialidase activity is defined based on enzyme unit (U), where one U is the amount of sialidase required to catalyse 1 nmol of BCIN per minute in a ml. The amount of sialidase activity corresponded to colour changes in the BV test bottle with ≥7.8U/ml for a positive result (blue or green) and 0 – 7.8U/ml for a negative result (yellow). The samples were defined as having high or low sialidase activity based on ≥7.8U/ml or 0 – 7.8U/ml, respectively (25).

### Leukocyte wet mount

Wet smears were prepared as previously described (62). Leukocyte counts were quantified using wet smears as leukocyte numerical range by observation of 10 nonadjacent microscopic fields and was used as a proxy for leukocyte activation at time of vaginal swab collection. Women were classified as having either a positive or a negative leukocyte result based on >15 or <15 counts/Hp of leukocytes (45, 63).

### Bioinformatics and statistical analyses

The raw sequencing reads were processed using the DADA2 package (v1.6.0) in R (v3.4.3)(64). The raw paired-end reads were assigned to samples based on their unique barcodes and truncated by cutting off the barcodes and primer sequences. The reads were then processed using the DADA2 package (v1.6.0) in R (v3.4.3) according to the following steps: quality controls, dereplication, error rate calculations, sequences denoising, paired-end reads merging, amplicon sequence variant (ASV) table constructing and removal of chimeras. The sequences in the ASV table were annotated using the RDP Classifier (65) with the GreenGenes database (66). The sequences assigned to *Lactobacillus* were further classified into species level using UCLUST (67) with the customised *Lactobacillus* database (V4 region extracted from representative sequences of *Lactobacillus* species from RDP database).

All downstream bioinformatic and statistical analyses were performed in R (v4.0.0)(68) unless otherwise stated. Low abundance taxa accounting for less than 0.1% within-sample relative abundance and present in less than 1% of samples were filtered using phyloseq (v1.32.0)(69) and genefilter (v1.70.0)(70) packages. This left a total of 167 ASVs representing 94 unique taxa. Extraction kit and reagent negative controls (n=103) were used to identify potential contaminants using the decontam package (v1.9.0) with a prevalence threshold of 0.1 and 0.5 respectively (71). A total of 12 contaminants were assigned as *Pseudomonas veronii*, *Bifidobacterium adolescentis*, *Klebsiella* spp., *Enhydrobacter* spp., *Chryseobacterium* spp., *Pedobacter* spp., *Bradyrhizobium* spp., *Methylobacterium* spp., *Escherichia coli*, *Acinetobacter johnsonii* and *Sneathia* spp., and *Species*_Other. Samples with a library size of <2000 reads were excluded from the analysis (n=90)(Fig. S1).

Each vaginal bacterial profile in the cohort was initially classified on the basis of the highest relative abundance of the dominant species taxon within the sample (2, 72). Dominant species with <30% abundance were classified as “Other” (n=16) and a single sample dominated by *Mesorhizobium* spp. was classified as “Outlier”. These samples were excluded from subsequent analyses. A total of 19 dominant (>30% relative abundance) species groups were then classified into VMG consisting of VMG I (*L. crispatus*); II (*L. gasseri*); III (*L. iners*); IV-A (BV-associated species; *Atopobium vaginae*, *Gardnerella* spp., *Megasphaera* spp., *Prevotella* spp., *Shuttleworthia* spp., *Veillonella* spp.)(36, 73); IV-B (Pathobionts; *Aerococcus* spp., *Anaerococcus* spp., *Streptococcus agalactiae*, *Streptococcus anginosus*, *Streptococcus luteciae*, *Streptococcus* spp.)(74); V (*L. jensenii*); VI (Other *Lactobacillus*; *L. delbrueckii*, *Lactobacillus* spp.) and VII (*Bifidobacterium* spp.). Analyses were also performed on samples classified into *Lactobacillus* species dominated and depleted groups based on *Lactobacillus* abundance of greater or lower than 90% (75).

Data were visualised using ggplot2 package (v3.3.1)(76). Statistical difference for continuous variables between groups were determined using either Mann-Whitney Test with Bonferroni p-adjusted values or Kruskal-Wallis Test with pairwise comparison between groups and Bonferroni p-adjusted values. Association between categorical groups were determined using Fisher’s Exact Test. For all statistical tests, significance was based on p<0.05 unless otherwise stated. Spearman’s correlation plots were generated from relative abundance using corrplot package (v0.84) with ward.D2 agglomerative method (77). To visualise diversity and richness for dominant species groups and individual samples from sub-cohort analysis, the Shannon diversity index was calculated using the phyloseq package (v1.32.0)(69) and richness was based on number of species observed. LefSe analysis was performed using per-sample normalisation to sum of values to 1M with a Linear Discriminant Analysis (LDA) threshold of 2.0 for discriminative features and an alpha of 0.05 for factorial Kruskal-Wallis test among classes and alpha 0.05 for pairwise Wilcoxon test between subclasses (78). Confusion matrices were computed using the caret package (v6.0-86) with a prevalence of 0.15 (79). Multiple logistic regression was performed using gestation at delivery (days) as a binary outcome (Preterm, <259 days and Term, ≥259 days) for the response variable and *Lactobacillus* abundance, sialidase activity and leukocyte wet mount results as explanatory variables.

### Data availability

Key analysis code and processed datasets required to reproduce the statistical analyses presented in this study are available the GitHub repository https://github.com/sherrianne. Raw data files for the sequence data used in this study are publicly available through the European Nucleotide Archive (https://www.ebi.ac.uk/ena) under accession numbers (PRJNA706523).

## Acknowledgements

We would like to thank all patients who have participated in this study. This work was funded by National Natural Science Foundation of China (NSFC81925062, 82002201), National Key R&D Program of China (2019YFA0802300) and the March of Dimes European Preterm Birth Research Centre at Imperial College London.

SN, MC, DAM and HZ conceived and designed the study. Clinical sampling and coordination of metadata was performed by MC, XW, ZDZ, HS, YH and YW. Experiments were performed by MC, ZY and WQ. Data processing and analysis was performed SN, MC and DAM. SN and MC prepared all figures and tables. SN and DAM wrote the first draft of the manuscript. All authors critically reviewed, read and approved the final manuscript.

## Supplementary Materials

Fig. S1. Study design (a) Summary of women included in final cohort analysis comparing women with and without outcome data and sub-cohort analysis of women with available maternal and neonatal data. (b) Density plot showing sample gestation of women with (red) and without (blue) outcome data segmented by trimester.

Fig S2. Vaginal microbiome groups of women without outcome data. Radial plot showing vaginal microbiome groups (VMG) of women without outcome data. Species classified into VMG I (*L. crispatus* dominated, blue); II (*L. gasseri* dominated, green); III (*L. iners* dominated, orange); IV-A (BV-associated, red); IV-B (pathobionts, pink); V (*L. jensenii* dominated, purple); VI (Other *Lactobacillus*, light blue) and VII (*Bifodobacterium* dominated, light purple). Outcome data was defined as availability of maternal age and birth gestation.

Fig S3. Lactobacillus dominant microbiome profiles of women with high and low sialidase activity. The microbiome profiles of women with *Lactobacillus* dominance and high or low sialidase activity dominated by Vaginal Microbiome Group (VMG) I (*L. crispatus*, blue), II (*L. gasseri*, green), III (*L. iners*, orange) and/or V (*L. jensenii*, purple).

Fig. S4. Microbiome composition and leukocyte presence or absence. Proportions of *Lactobacillus* abundance, vaginal microbiome groups (VMG), sialidase activity and leukocyte wet mounts based on chorioamnionitis presence (Chorio+) and absence (Chorio-).

Table S1. Relative abundance and Shannon diversity index of dominant species groups.

Table S2. Vaginal microbiome groups (VMG) of women with and without outcome data.

Table S3. Relative abundance of *Lactobacillus*, *Bifidobacterium* and BV-type species in women with high or low sialidase activity.

Table S4. Birth gestation of women based on *Lactobacillus* abundance, sialidase activity and leukocyte wet mount results.

Data set S1. Metadata and associated taxanomic assignments.

